# Tropomyosin isoforms segregate into distinct clusters on single actin filaments

**DOI:** 10.1101/2021.07.23.453476

**Authors:** Peyman Obeidy, Thomas Sobey, Philip R. Nicovich, Adelle C. F. Coster, Elvis Pandzic

## Abstract

Tropomyosins (Tpm) are rod-shaped proteins that interact head-to-tail to form a continuous polymer along both sides of most cellular actin filaments. Head-to-tail interaction between adjacent Tpm molecules and the formation of an overlap complex between them leads to the assembly of actin filaments with one type of Tpm isoform in time and space. Variations in the affinity of tropomyosin isoforms for different actin structures are proposed as a potential sorting mechanism. However, the detailed mechanisms of spatio-temporal sorting of Tpms remain elusive. In this study, we investigated the early intermediates during actin-tropomyosin filament assembly, using skeletal/cardiac Tpm isoform (Tpm1.1) and a cytoskeletal isoform (Tpm1.6) that differ only in the last 27 amino acids. We investigated how the muscle isoform Tpm1.1 and the cytoskeletal isoform Tpm1.6 nucleate domains on the actin filament and tested whether (1) recruitment is affected by the actin isoform (muscle vs cytoskeletal) and (2) whether there is specificity in recruiting the same isoform to a domain at these early stages. To address these questions, actin filaments were exposed to low concentrations of fluorescent tropomyosins in solution. The filaments were immobilized onto glass coverslips and the pattern of decoration was visualized by TIRF microscopy. We show that at the early assembly stage, tropomyosins formed multiple distinct fluorescent domains (here termed “cluster“) on the actin filaments. An automated image analysis algorithm was developed and validated to identify clusters and estimate the number of tropomyosins in each cluster. The analysis showed that tropomyosin isoform sorting onto an actin filament is unlikely to be driven by a preference for nucleating on the corresponding muscle or cytoskeletal actin isoforms but rather is facilitated by a higher probability of incorporating the same tropomyosin isoforms into an early assembly intermediate. We showed that the 27 amino acids at the end of each tropomyosin seem to provide enough molecular information for attachment of the same tropomyosin isoforms adjacent to each other on an actin filament. This results in the formation of homogeneous clusters composed of the same isoform rather than clusters with mixed isoforms.

## Introduction

Tropomyosins (Tpms) form co-polymers with actin and are composed of a family of over 40 isoforms. Cytoskeletal Tpms appear largely form homodimers which in turn form homopolymers with actin and define the compositional variety of actin filaments (Gunning *et al*, 2015). Their interaction with actin leads to specialization of actin filament function in time and space (Gimona, Watakabe, and Helfman, 1995; Gunning, O’Neill and Hardeman, 2008). Due to alternative splicing, most tropomyosin molecules have differentially encoded N- and C-termini. The overlap region plays an important role in determining the orientation and dynamics of tropomyosin dimers on filamentous actin (Sliwińska and Moraczewska, 2013). Multiple studies have established that the N- and C-terminal overlapped regions are not functionally equivalent (Moraczewska, Nicholson-Flynn, and Hitchcock-Degregori, 1999). Differences in these sequences affect the binding of tropomyosin to actin as well as that of other actin-binding proteins (Mak and Smillie, 1981; Hammell and Hitchcock-DeGregori, 1996). The current Tpm/F-actin binding model supports tropomyosin molecules interacting head-to-tail to form a homogeneous continuous coiled-coil structure floating in the positively charged major groove of an actin filament (Gunning, O’Neill and Hardeman, 2008; von der Ecken *et al.*, 2015). This assembly model also postulates that cooperativity in tropomyosin binding is driven by an 8-11 amino-acid overlap between the amino and carboxyl termini of adjacent tropomyosin dimers (Tobacman, 2008). Although it has been shown that the overlap regions could lead to distinct functions for Tpms (Moraczewska, J. *et al.*, 1999), the detailed mechanism of this interaction remains elusive. To elucidate the intrinsic mechanism of tropomyosin sorting to the actin filament, we employed a single-molecule imaging approach based on total internal reflection fluorescence (TIRF) microscopy and microfluidics. We investigate whether the amino acid differences between the labeled Tpms are sufficient to guide and discriminate between these isoforms during the early stages of tropomyosin assembly. We selected the tropomyosin dimers, Tpm1.1 and Tpm1.6, which are identical in their amino-acid sequences apart from their C-termini (last 27 amino acids) that are encoded by different exons (Gunning, O’Neill and Hardeman, 2008).

In a typical TIRF setup, light is reflected at an interface between two media with different refractive indices. It generates an evanescent field that effectively produces a ~150 nm thick exponentially decaying excitation profile (Axelrod *et al.* 1984). Although restricting excitation to a very thin section near the coverslip allows single-molecule detection, a lack of automated tools for quantification and resolving the number of molecules within sub-diffraction limited clusters remains challenging. Here we combine single-molecule detection with the use of microfluidic devices to facilitate and automate image processing to investigate the early assembly of tropomyosins bound to actin filaments. We deconvolved and counted the number of tropomyosins in bright diffraction-limited clusters based on photobleaching (Zhang, H. and Guo, P., 2014) and intensity-distribution-based deconvolving approaches (Mutch, S. A. *et al.* (2007); Peterson, E. M. and Harris, J. M., 2010).

In this paper we report that segregation of Tpm isoforms may be intrinsic to the different proteins. We have visualized and quantified snapshots of the assembly process by reconstituting actin filament in the presence of labeled cytoskeletal (Tpm1.6) and skeletal (Tpm1.1) Tpm isoforms which bind to different sites on actin filaments. We used the closely related recombinant tropomyosin isoforms from the TPM1 gene, skeletal-muscle Tpm1.1 and cytoskeletal Tpm1.6 isoforms. Our findings show that at low concentrations, Tpms (1.1 and 1.6) bind multiple locations along naked *α*- and *β*-actin filaments to form small clusters. No intrinsic preferences were observed for sorting of the correct Tpm to its biologically relevant actin filament; for example, Tpm1.6 binds with higher affinity than Tpm1.1 to both actin isoforms.

## Materials and Methods

### Expression, purification, and labeling of Tpms

Expression of recombinant human Tpm1.6 (Tm2) and rat Tpm1.1 (αTm) with N-terminal Ala-Ser extension in *E. coli* BL21 (DE3) cells using pET3a+ expression vectors was carried out as described previously (PMID: 8144630, Monteiro PB, JBC 1994). The protein was purified using ammonium sulphate fractionation, dialysis, chromatography, and gel filtration chromatography and then lyophilized for storage at −20 °C as described previously (Janco M. et al 2016). The lyophilized protein was resuspended in 1-2 mL T buffer (150 mM NaCl, 10 mM Tris-HCl pH 7.5, 2 mM MgCl2) supplemented with 5 mM DTT and then dialyzed against T buffer containing 0.5 mM DTT (2×2 h). The Cys-190 residues of the Tpm1.6 and Tpm1.1 dimers are prone to form a disulphide. Thus, the protein was reduced prior to labeling by incubation in the presence of 20 mM DTT at 37°C for 2 h. To remove the DTT, the buffer was then exchanged with T buffer using desalting columns (Zeba Microspin, 7k MWCO, Thermo Fisher Scientific). Tpms were then reacted at Cys-190 with a five-fold molar excess of Alexa Fluor 488-C5-maleimide (Ex/Em:495/519 nm) or Alexa Fluor 568-C5-maleimide (Ex/Em:578/603 nm) (Life Technologies) in the dark at 37°C for 2 h. The reaction was quenched using β-mercaptoethanol added at a two-fold molar excess over the dye. Finally, the unconjugated dye was removed using Zeba columns (up to five times). The purity of the labeled mixture was examined using SDS-PAGE with fluorescence detection (UV excitation) and Coomassie Blue staining. The labeled protein concentration was measured using the Direct Detect spectrometer (Merck Millipore). The degree of labeling was calculated as the ratio of the dye concentration and the protein concentration, whereby the dye concentration was determined from the absorption of the sample at the wavelength corresponding to the maximum absorption for the dye and the extinction coefficient.

### Co-sedimentation of tropomyosin-actin

The affinities of unlabeled and labeled tropomyosins for actin filaments were measured using a standard co-sedimentation assay (Heald, R. *et al* 1988). Prior to co-sedimentation, tropomyosin isoforms were incubated with 1 mM DTT and heated in a water bath at 56°C for 5 min. Increasing concentrations of tropomyosins (0.2-3.5 μM) were mixed and incubated with 3 μM rabbit skeletal-muscle F-actin (Cytoskeleton, Inc.) in T buffer at a final reaction volume of 50 μL for 1 h at room temperature. Following the incubation, the samples were spun in a bench-top Beckman ultracentrifuge at 670,000 g at 20 °C for 30 min (TLA-120.1 rotor, Beckman-Coulter) to pellet F-actin and associated Tpm. Supernatant and pellet fractions were resolved on a 12% polyacrylamide gel, and proteins were visualized by Coomassie Blue staining and the protein bands were quantified by densitometry (Epson Perfection V750 Pro scanner and ImageJ). The ratio of the Tpm/actin density was plotted as a function of the concentration of free Tpm in the supernatant determined by densitometry. The Hill equation (Equation (1)) was used to fit the binding curves:

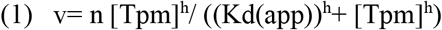

where v is the Tpm/actin density ratio, [Tpm] is the concentration of free Tpm, n is the maximum Tpm/actin density ratio, h is the Hill coefficient and Kd(app) is the apparent equilibrium dissociation constant for Tpm binding to F-actin (Janco M. et al 2016).

### Single-molecule photobleaching

Freshly cleaned coverslips were clamped into a magnetic chamber (Chamlide CMB, Live Cell Instruments), and labeled tropomyosins (Tpm1.6 AF488, Tpm1.6 AF568, Tpm1.1 AF488, and Tpm1.1 AF568) were sparsely adsorbed to the coverslips from solution (32 pM) in TIR50 buffer (50 mM NaCl,10 mM Tris-HCl pH 7.8, 2 mM MgCl_2_, 1 mM ATP, 0.2 mM CaCl_2_ and 50 mM DTT) supplemented with additional NaCl to a final concentration of 150 mM to prevent oligomerization. After incubation (5 min) the surface was rinsed and the solution in the chamber was replaced with 200 *μ*L of TIR50 buffer.

Tropomyosin dimers appeared as a diffraction-limited spot in the time-lapse images collected using Total Internal Reflection Florescence (TIRF) microscopy. Images were collected at an imaging rate of 5 frames per second until most of the spots were photobleached. Spots with two bleaching steps (corresponding to the two fluorophores on a tropomyosin dimer) were selected and fitted with a 2D Gaussian function to determine the amplitude and half-width (sigma, σ) of the diffraction-limited signals. Mean values for amplitude and σ were determined by fitting the corresponding histograms of the data (~100 photobleaching events) with a Gaussian distribution.

### Preparation of microfluidic flow cells

Coverslips (Marienfeld superior No. 1.5H, 24 mm × 60 mm) were sonicated in acetone for 5 min followed by rinsing with Milli-Q water. Filtered N2 gas was blown over the coverslip before passing the coverslip briefly through the flame of a Bunsen burner. The dried coverslip was treated in a plasma cleaner for 3 min at a pressure of ~650 Torr. Microfluidic devices with channel dimensions of 1×0.8×0.04 mm (*L*×*W*×*H*) were prepared from polydimethylsiloxane (PDMS) using replica molding and cured in an oven at 70°C for 2 h. The PDMS device was punctured at the channel ends with a biopsy punch (1 mm diameter) for connecting the device to tubing and the device was then cleaned with isopropanol. The PDMS device was placed onto a freshly plasma-cleaned coverslip and adhered together by placing the assembled device under a weight (600 g) for 3 min. Finally, polyethylene tubing (PE20 Intramedic, Clay Adams, the internal diameter of 0.38 mm) was inserted into the holes at the inlet and outlet of the channels in the assembled device.

### The capture of actin filaments on modified coverslip surfaces

The microfluidic channels were filled with a solution of a copolymer of poly-l-lysine (PLL) and poly (ethylene glycol) (PEG) [Susos AG, PLL (20)-g[3.4]-PEG (2)/PEG (3.4)-biotin (20%)] in PBS (5 μL, 1 mg mL^−1^) and the copolymer was allowed to adsorb to the glass surface for 20 min at room temperature. The unbound PLL-PEG was washed out of the channel with 20 *μ*L of PBS. The coated surface was then treated with blocking buffer (20 mM Tris pH 7.5, 2 mM EDTA, 50 mM NaCl, 0.03% NaN3, 0.025% Tween 20, 0.2 mg/mL BSA) at room temperature in a humid chamber for 10 min. The outlet tubing of the PDMS device was connected to a syringe pump operated in withdraw mode, and each channel was subsequently rinsed with 20 μL of PBS followed by 20 μL of 5% BSA, incubated for 10 min and subsequently washed with 20 *μ*L of D buffer (10 mM Tris-HCl pH 7.8, 2 mM MgCl2, 1 mM ATP, 0.2 mM CaCl2 and 50 mM DTT). This surface was used for non-specific capture of actin filaments decorated with labeled tropomyosin. Labeled tropomyosins (0.015 *μ*M of each isoform) were mixed prior to adding 1 *μ*M monomeric actin (G-actin, Hypermol) in D buffer. Actin polymerization was triggered by adding 50 mM NaCl and incubating the mixture for 5-10 min. Subsequently, the reaction mixture was gently mixed with a solution of polyethylene glycol (PEG 300) to a final concentration of 10% v/v, just before injecting 3-6 *μ*L of the mixture into the microfluidic chambers. The channel was then rinsed with TIR50 buffer at a flow rate of 5 *μ*L/min. Decorated filaments were allowed to bind to the surface and unbound filaments were removed by washing the microfluidic channel with TIR50 buffer at a flow rate of 5 *μ*L/min with 20 *μ*L.

### TIRF Imaging of labeled Tpm1.1 and Tpm1.6 binding to actin filament

Single-molecule experiments were conducted using an inverted TIRF microscope (TILL Photonics) with Andor iXon3 897 Ultra back-illuminated EM-CCD cameras (Hamamatsu) and 100X/1.46 Alpha-Plan apochromatic Oil 0.17mm (UV) Vis-IR objective. Images were acquired using the TILL Photonics microscopy software packages Live Acquisition (LA) and Offline Analysis (OA). Dual-color time-lapse TIRF imaging was conducted using 488 nm (Dynamic Laser, Salt Lake City, UT) and 568 nm (Coherent, Santa Clara, CA) lasers at 20% (~1 mW) laser power, a multiplication gain setting of 300, 100 ms laser exposure time and an imaging frequency of 5 Hz.

### Calibrating tropomyosin numbers on an actin filament

Clusters of tropomyosin dimers bound to actin filaments appeared as punctate signals in the TIRF images. Tropomyosin clusters were fitted with 2D Gaussian function and then iteratively deconvolved from the overlapping clusters, until the background level is reached. The number of fluorophores in each cluster was then calculated by dividing the total integral intensity of the diffraction-limited spot of a cluster by that of a single fluorophore. For simplicity, we will refer to the total integral intensity of a diffraction-limited spot as a volume. An automated MATLAB algorithm (see supplementary material) was used to fit clusters on each filament and calculate the total number of fluorophores per filament. The algorithm was validated on datasets with filaments with a sparse pattern of labeled tropomyosins and using simulated cluster data. The algorithm was designed for processing a large number of filaments from independent experiments and distributions of cluster sizes (number of fluorophores per cluster) and the distances between clusters to be determined. The following steps were taken to correct these misidentifications: (a) peaks within a distance of 2*σ* pixels of each other, equivalent to the standard deviation, *σ,* of the point-spread function (PSF), (b) small intensity peaks (with *σ* <1) remaining after this process were assigned to noise, as it is physically impossible to have PSF with *σ* <1 pixel, and excluded from further analysis (c) peaks with intensity value equivalent to a *σ/2* or smaller and more than three pixels were excluded, (d) filaments with only one type of tropomyosin (usually small filament) or only one of each tropomyosin were also excluded.

## Results

### Labeled tropomyosins bind cooperatively to actin filaments

Both Tpm1.1 and Tpm1.6 isoforms, derived from the TPM1 gene, were labeled on their only cysteine at position 190 using maleimide derivatives of Alexa Fluor 488 (AF488) and 568 (AF568) (Fig. 1A and 1B). These Tpm isoforms contain a few aromatic amino acids and have a relatively low extinction coefficient, 5960 and 8940 cm^−1^M^−1,^ respectively. Thus, we calculated the degree of labeling ratio by dividing the concentration of the fluorophore (measured using UV light absorption) in labeled protein by the concentration of the protein measured using (infrared (IR) absorption). The monomer labeling ratio was determined to be approximately 67 and 95% for Tpm1.1 and Tpm1.6, respectively (Fig. 1C). Assuming a random incorporation of the dye, this labeling ratio corresponds to a mixture containing Tpm dimers labeled with two, one or no fluorophores, whereby singly labeled dimers constitute the main fraction. Subsequently, we evaluated the impact of the labeling process on the binding of labeled Tpm1.1 and Tpm1.6 to skeletal-muscle F-actin using the standard co-sedimentation assay (Fig. 1D, 1E). Our findings show that Alexa Flour 488 (AF488) and 568 (AF568) maleimide labeling of Tpm1.1 and Tpm1.6 did not affect the formation of Tpm-actin filaments, nor was the cooperative nature of the binding affected (Table 1). These results agree with previously reported studies of the retention of the binding ability of Tpm1.1 containing modified 190 cysteine residues (Betcher-Lange and Lehrer, 1978; Ishii and Lehrer, 1986).

**Table 1.** Binding properties of unlabeled and labeled tropomyosin isoforms. Data are mean±S.E. combined from 2-3 independent experiments (n=1, Tmp1.1 Alexa Fluor 568).

| Tm | K<sub>dapp</sub> (μM) | α<sup>H</sup> |
| --- | --- | --- |
| Tpm | 0.64 ± 0.06 | 3.93 ± 1.06 |
| αTm AF488 | 0.17 ± 0.02 | 3.52 ± 1.39 |
| αTm AF568 | 0.24 ± 0.06 | 3.44 ± 3.67 |
| Tm2 | 0.44 ± 0.07 | 1.86 ± 0.54 |
| Tm2 AF488 | 1.21 ± 0.17 | 2.36 ± 0.59 |
| Tm2 AF568 | 2.37 ± 1.38 | 2.03 ± 0.83 |

**Figure 1.**
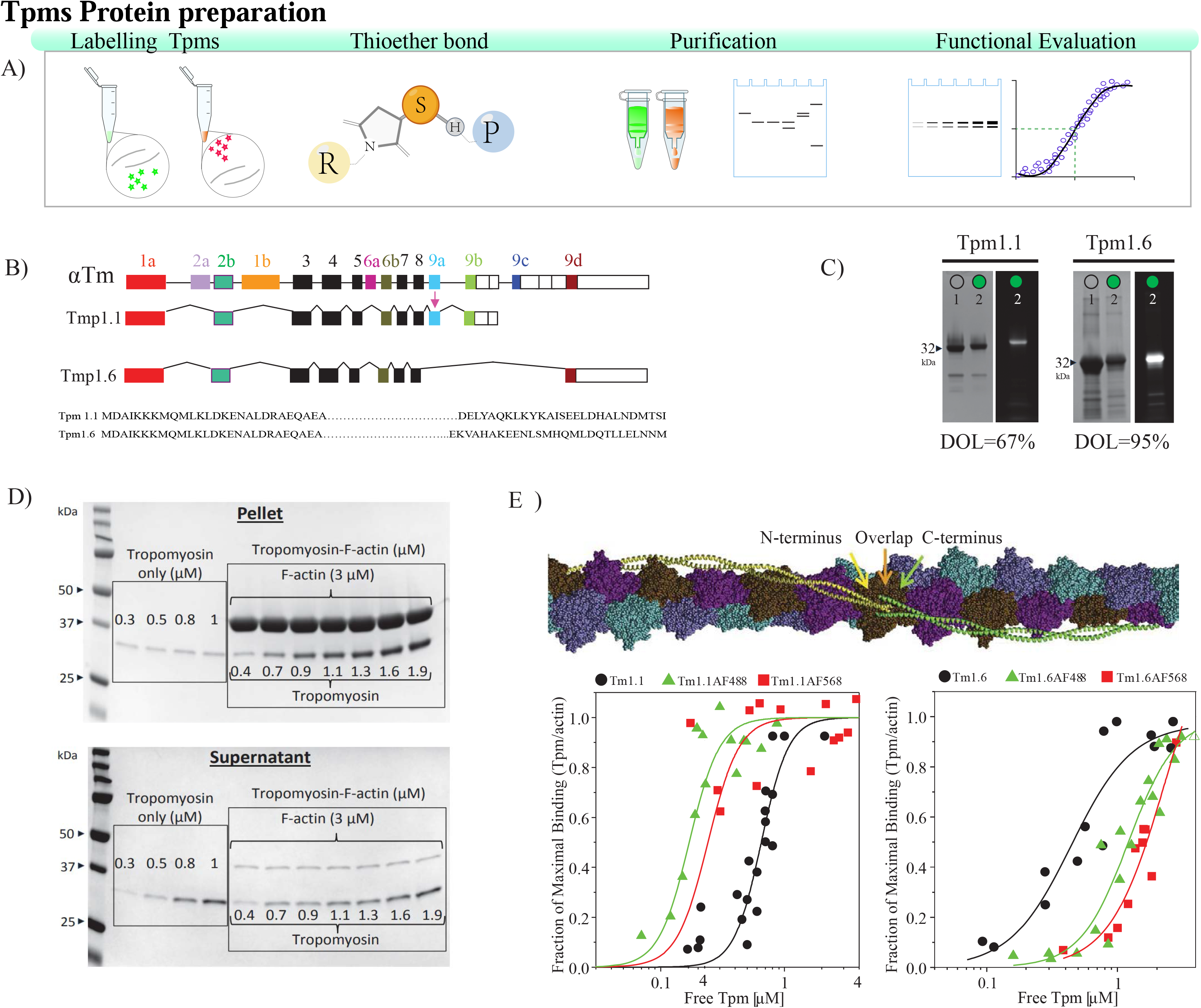
Labeled tropomyosins bind cooperatively to actin filaments. (A) Schematic illustration of Tpms preparation, labeling and functional evaluation. (B) Diagram of Tpm1.1 (αTm) and Tpm1.6 (Tm2) isoforms. (C) Cysteine residues at position 190 on the recombinant tropomyosin dimers were labeled with Alexa Fluor 488 (ɛ = 71000) or 568 (ɛ = 91300) C5 maleimide. Image adapted from [164]. (C) Representative SDS-PAGE with Coomassie Blue staining (left) and fluorescence detection (right). Lane 1, unlabeled protein; lane 2, a protein labeled with Alexa Fluor 488 C5 maleimide dye. (D) SDS-PAGE gels of the (left panel) supernatant and (right panel) pellet fractions from sedimentation of Tpms with filamentous α-actin stained with Coomassie Blue. (E) Tpms interact head-to-tail on the actin filament. (F) Binding curves were obtained from co-sedimentation of unlabeled and labeled (right panel) Tpm1.1 and (left panel) Tpm1.6 to skeletal-muscle F-actin. Data points were combined from three independent experiments except for Tpm1.6 n=2. The B and E images were adapted from Gunning *et al.* 2005, Orzechowski *et al.* (2014), respectively.

### Single-Molecule Calibration Based on Photobleaching

To calibrate the properties of single labeled Tpms, we collected the data for the diffraction-limited spots with one- and two-step photobleaching profiles. The two-step photobleaching profile was broken into two single-step profiles before pooling the values. Photobleaching traces were generated by fitting the fluorescent spot in each frame was with a 2D symmetric Gaussian function and calculating its volume using equation (1);

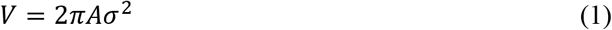

where A represents the amplitude (height of the Gaussian function above the background intensity) and σ is the standard deviation. The intensity of a single fluorophore was then determined as the step height between intensity levels in the photobleaching profiles. The mean intensity and standard deviation of a single fluorophore were determined by fitting the distribution of step heights with a Gaussian function (Fig. 2D). The data is summarized in Table 2. We further determined the number of fluorophores for each spot present in the first frame of the fluorescence movie used initially for the photobleaching analysis. The volume of the spot was divided by the volume of a single fluorophore. The resulting distribution confirmed that 81% of the Tpm dimers were labeled with a single fluorophore (data not shown). This data was also consistent with the aforementioned labeling ratio determined by spectroscopy.

**Table 2.** Photobleaching properties of labeled tropomyosin dimers. Note that units for σ are in pixels. Both Amplitude (Amp) and integral values are in arbitrary units.

| Single Molecule | Bleach Profile | Mean( σ) | Mean(Amp) | $\frac{\text{First Step}}{\text{Second Step}}$ | Integral |
| --- | --- | --- | --- | --- | --- |
| αTm AF568<br>(n=94) | First Step | 1.56 ± 0.16 | 544± 217 | 0.87 | 10584± 10108 |
|  | Second Step | 1.55 ± 0.14 | 614± 446 |  | 11049± 4427 |
|  | Combined | 1.55± 0.16 | 603± 395 |  | 10171± 6855 |
| αTm AF488<br>(n=149) | First Step | 1.37 ± 0.15 | 433± 218 | 1.06 | 5418± 6633 |
|  | Second Step | 1.36 ± 0.13 | 409± 267 |  | 5677± 2209 |
|  | Combined | 1.37± 0.16 | 426± 283 |  | 5182± 4061 |
| Tm2AF568<br>(n=66) | First Step | 1.55 ± 0.25 | 487± 245 | 0.87 | 9579± 14245 |
|  | Second Step | 1.60 ± 0.26 | 560± 787 |  | 13358± 5036 |
|  | Combined | 1.58± 0.23 | 642± 607 |  | 11719± 9031 |
| Tm2AF488<br>(n=115) | First Step | 1.34 ± 0.13 | 427± 208 | 0.79 | 6066± 8451 |
|  | Second Step | 1.35 ± 0.12 | 538± 285 |  | 6787± 2587 |
|  | Combined | 1.35± 0.13 | 482± 339 |  | 6341± 4538 |

**Figure 2.**
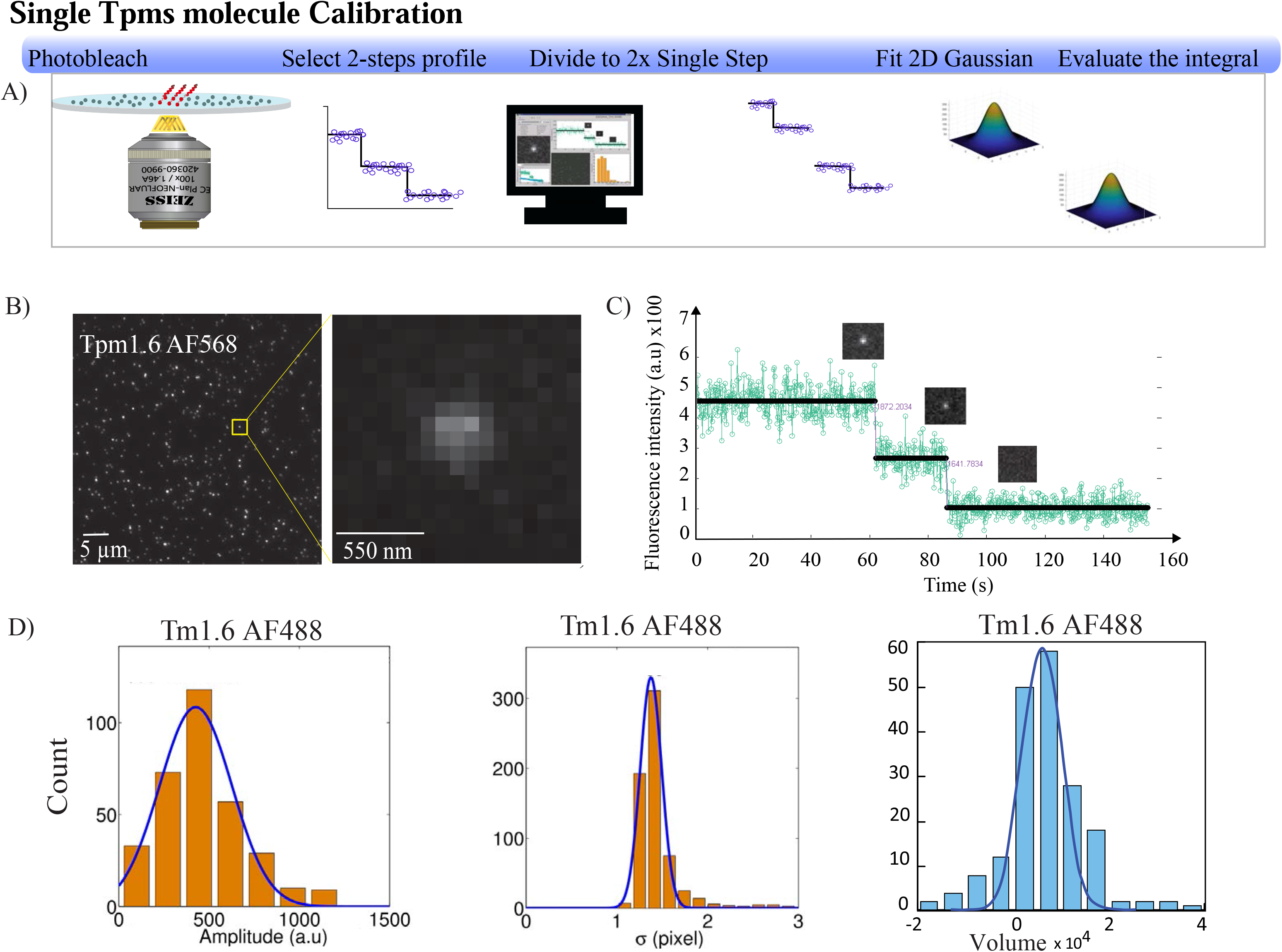
Single-Molecule intensity calibration Based on Photobleaching. Schematic illustration of Tpms calibration as a single molecule. (B) Representative TIRF image of labeled tropomyosin adhered to the coverslip. Each labeled molecule appears as a diffraction-limited spot. (C) Stepwise photobleaching of a diffraction-limited spot as a function of time. (D) Distributions of the fluorescence amplitudes (left panel) Standard deviation (σ) (center panel) and total integral (volume) of detected spot (right panel) for the fitted Gaussian profiles of the single-molecule (see Table 2).

### Observation and quantification of actin-tropomyosin filament assembly at the single-molecule level

To visualize and evaluate the binding patterns of the tropomyosins on actin filaments, we used custom-made microfluidic devices (Fig 3A-C). The mixture containing either α- or β-actin monomers and labeled Tpms was incubated for 5 minutes and actin polymerization was initiated by increasing the ionic strength of the mixture using NaCl salt (50 mM final concentration). The binding of the labeled Tpms to actin filaments was enhanced using the crowding effect of 10% PEG 300. The decorated filaments were non-specifically captured on the surface of the glass coverslip modified with PLL-PEG and 5% BSA and imaged using TIRF illumination (Fig 3C). We divided experiments into different conditions: binding of the same Tpm isoform labeled with either AF488 or AF568 (“control”) and binding of varying Tpm isoforms, each labeled with a different fluorophore (“competition”) (Fig. 3C and 3D). The bound tropomyosin appeared as a diffraction-limited spot in all experimental conditions, which we call “clusters”. The images were further analyzed using the algorithm developed for this study.

**Figure 3.**
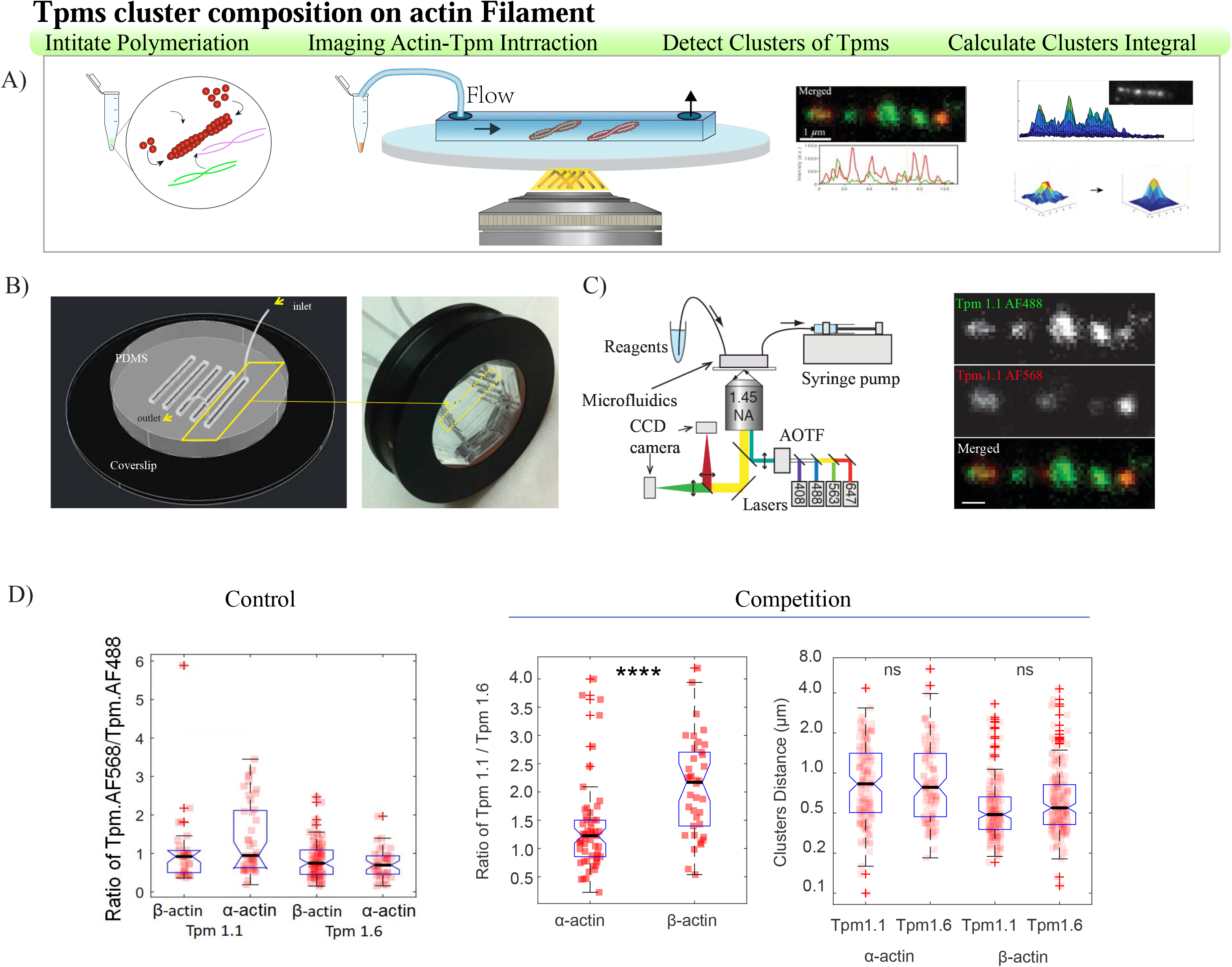
Observation and quantification of tropomyosin molecules bound to actin filaments assembly at the single-molecule level using a TIRF microscope configuration and flow-cell design. Schematic illustration of capturing decorated filaments to the surface. (B) Actin filaments are decorated with the fluorescent Tpm molecules in solution and then injected into the flow channel of the microfluidic imaging device for capture on the chemically modified coverslip. (B) PDMS device assembly. (C) Fluorophores captured on the filament are excited with laser light, which can be modulated with an acoustic-optical tunable filter (AOTF). The emission light is collected via a high-numerical-aperture objective and detected with a sensitive electron-multiplying charge-coupled-device (CCD) camera. The emission path includes a beam splitter for simultaneous dual-color imaging (left panel). The representative images of actin filaments are decorated with labeled tropomyosins (right panel). (D) The distribution of the same tropomyosin isoforms’ ratio is labeled with two different fluorophores on actin filaments (control experiments). Actin (1 μM) was co-assembled with tropomyosin (0.015 μM of each label) prior to the capture of the decorated filament (right panel). The distribution of the ratio of the different tropomyosin isoforms labeled with two different fluorophores are shown on actin filaments (competition experiments, mid-panel). Data pooled for Tpm1.1AF488 and Tpm1.1.AF568 into Tpm1.1 and similarly for Tmp1.6. The distribution between consecutive clusters on actin filaments in a competition experiment (left panel). Data are pooled from three independent and represented as mean±s.d., t-test, ****P < 0.0001. ns denotes not significant.

Control experiments showed that the AF568-labeled Tpm and AF488-labeled Tpm were equally incorporated into clusters on the actin filament, whereas Fig 4D (first panel) shows a slight increase in the occupation ratio of Tpm1.1 AF568 over the same isoform labeled with AF488 for α-actin. This finding, in addition to the biochemical assays, further validated the absence of any adverse effect of the small fluorophores AF488 and AF568 on the affinity of tropomyosins for actin. Thus, the tropomyosins with fluorophore are referred to as “labeled tropomyosin” in the “competition experiment”. Nevertheless, the “competition” experiments showed an average of 1.2, and 2.2 times more labeled Tpm1.1 than Labeled Tpm1.6 incorporated into clusters on α-actin and β-actin filament, respectively. This suggests that, at low concentrations, Tpm1.1 exhibits a higher occupation fraction for both β-actin and α-actin, indicating that it shows a higher affinity than Tpm1.6 to both actin isoforms. This is in agreement with previous studies using biochemical assays (Skórzewski, Śliwińska, *et al.*, 2009). (Fig. 3A-D).

**Figure 4.**
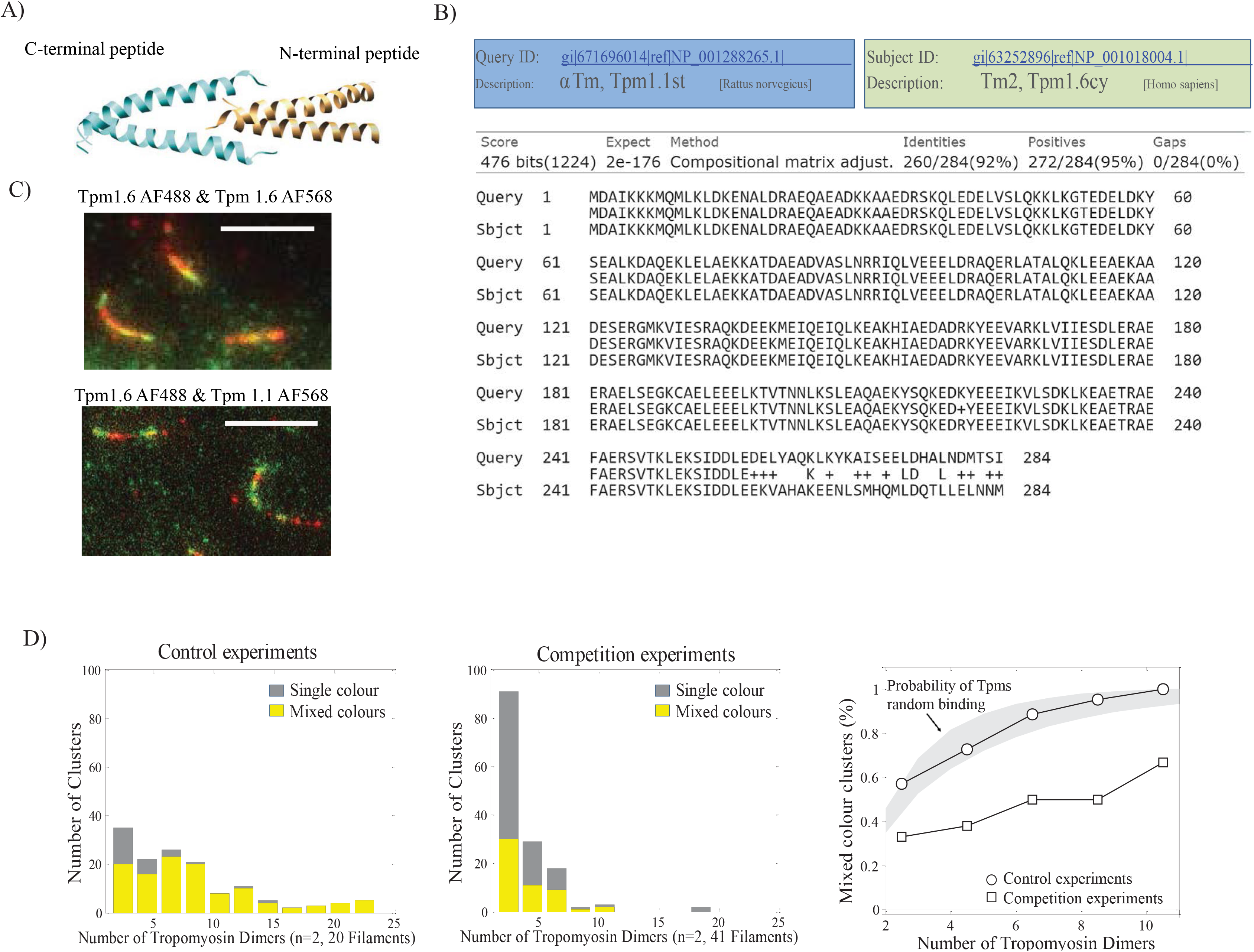
Different tropomyosin isoforms separate into distinct clusters. (A) An illustration of the NMR structure of the C-and N-terminal overlapped peptides with the 8-11 residues spans (adapted from Greenfield *et al.*, 2006). (B) Alignment of Tpm1.1 amino-acid sequence against Tpm1.6 sequence with the 27^th^ amino acid difference. (C) Representative TIRF images of similar tropomyosins mixed in the same clusters (yellow) clusters (top panel) vs different tropomyosin isoforms are separating into distinct clusters (bottom panel). (D) Histograms of the number of tropomyosin dimers per cluster for single-color and mixed-color clusters for control experiments (20 filaments from two independent experiments, right panel) and competition experiments (41 filaments from two independent experiments, mid-panel). The fraction of mixed-color clusters as a function of cluster size (number of tropomyosin dimers per cluster). The grey line shows the fraction of mixed clusters predicted for random incorporation of both colors based on a binomial distribution with the experimentally determined probabilities of incorporating red (labeled with AF568) and green (labeled with AF488) tropomyosins (left panel).

We also competed two the tropomyosins using both skeletal and cytoskeletal actin filaments and found that the distributions of inter-cluster distances were similar for all combinations of actin and tropomyosin isoforms. Notably, we observed more clusters with shorter distances between Tpm1.1s and Tpm1.6s consecutive clusters on β-actin Table 3D.

**Table 3.** Summary of quantification of labeled tropomyosins decorated single actin filaments. Data are pooled from three to four independent experiments and represented as (median, 25%, 75% quartiles, the total number of diffraction-limited spots).

| Tpm1.1 |  | Tpm1.6 |  |
| --- | --- | --- | --- |
| on β-actin | on α-actin | on β-actin | on α-actin |
| Ratio of Tpm in Ch1(red,AF568)/Ch2(green, AF488) |  |  |  |
| 0.9 (0.5, 1, n=33) | 0.9 (0.6, 2, n=44) | 0.7 (0.5, 1, n=79) | 0.7 (0.5, 0.9, n=39) |
| Filament Length (μm) | | | |
| 4 (3, 6, n=34) | 5 (4, 6, n=46) | 4 (4, 6, n=50) | 7 (5, 9, n=43) |
| Tpm number per cluster |  |  |  |
| 3 (2, 6, n=225) | 3 (2, 6, n=536) | 2 (1, 4, n=584) | 2 (1, 4, n=301) |
| Number of clusters per filament |  |  |  |
| 3 (2, 4, n=66) | 5 (3,7, n=90) | 3 (2, 5, n=158) | 3 (1, 4, n=84) |
| Cluster Distances (μm) | | | |
| 0.6 (0.5, 1, n=157) | 0.5 (0.4, 0.7, n=442) | 0.6 (0.4, 1, n=413) | 0.9 (0.5, 2, n=202) |

**Competition Experiments**
| Ratio of Tpm 1.1 vs Tpm 1.6 |  |  |  |
| --- | --- | --- | --- |
| α-actin | | β-actin | |
| 1.2 (0.9, 2, n=57) |  | 2.2 (1, 3, n=38) |  |
| Filament Length (μm) | | | |
| 5 (4, 6, n=64) |  | 5 (4, 6, n=43) |  |
| Tpm number per cluster |  |  |  |
| Tpm1.1 | Tpm1.6 | Tpm1.1 | Tpm1.6 |
| 2 (1, 4, n=229) | 2 (1,3, n=176) | 5 (2, 13, n=219) | 2 (1,5, n=229) |
| Number of clusters per filament |  |  |  |
| 4 (2, 5, n=62) | 2 (1, 4, n=62) | 5 (3,9, n=41) | 5 (4, 8, n=41) |
| Cluster Distances (μm) | | | |
| 0.8 (0.5, 1, n=146) | 0.8 (0.5, 1, n=115) | 0.49 (0.4, 0.7, n=179) | 0.55 (0.4, 0.8, n=189) |

### Different tropomyosin isoforms separate into distinct clusters

To investigate whether the C-termini (27 C-termini amino acids) differences were sufficient to discriminate between these isoforms during the early stages of tropomyosin assembly, we quantified the composition of clusters using the method presented above. By selecting the clusters containing at least two tropomyosin dimers (i.e. at least two fluorophores), we calculated the fraction of single-color and mixed-color clusters as a function of cluster size (Fig. 4D, right and middle panel). We observed that the percentage of mixed-color clusters increased with the number of tropomyosin dimers per cluster. To determine if the observed mix-color portions were due to a random probability of Tpms interaction with the actin filament, we evaluate the probability of the mixed incorporation of tropomyosins using a binomial distribution with the experimentally determined probabilities.

Our data shows that in control experiments (same isoform labeled with different colors), tropomyosin dimers bound independently to the actin filament and did not bind as aggregates. The curve determined for the control experiment was similar to the curve predicted for the random incorporation of both colors (Fig. 4D, left panel). In contrast, exposing actin filaments to different isoforms of labeled Tpm1.1 and Tpm1.6 led to a pattern of the decoration with a reduced percentage of mixed clusters compared to the control experiments (Fig. 4D, central panel). This observation suggests that the two isoforms are not randomly incorporated into a growing tropomyosin strand on an actin filament. There appears to be a higher probability that the same isoform rather than different isoforms will be incorporated into the cluster during assembly.

## Discussion

This study prepared and labeled equimolar concentrations of rat skeletal-muscle Tpm1.1 and cytoskeletal Tpm1.6 tropomyosin molecules. Using labeled protein, we decorated single filaments of muscle and cytoskeletal actin prior to capturing the filaments on the modified microfluidic chamber surface. Both isoforms were expressed from the same gene with only a 27 amino acid difference in their structure. An unexpected outcome of our measurements was despite the small differences in the structures, our results indicate that the difference was sufficient to cause preferential neighbouring binding of like isoforms.

Tropomyosin isoforms in their native environment get sorted into specific populations of actin filaments. Some isoforms differ in only one exon, and this difference is sufficient to localize them to a particular population of actin. For example, Tm5NM1 (Tpm3.1) has been mainly observed with stress fibers and Tm5NM2 (Tpm3.2) with actin filaments associated with the Golgi apparatus (Percival *et al.*, 2004). An essential determinant for tropomyosin binding, polymerization, and function are the overlapping complex between the N- and C-termini of adjacent tropomyosin dimers (Moraczewska, Nicholson-Flynn, and Hitchcock-Degregori, 1999; Masedunskas *et al.*, 2018). More recently, Tpm localization was also suggested to be the effect of the overlap region in an isoform-specific manner. However, differences obtained for Tpm1.1 and Tpm1.6 in intravital imaging were comparatively small to indicate any functional differences (Sliwińska and Moraczewska, 2013). Another study, using cross-linking of F-actin and fluorescence changes in F-actin labeled with acrylodan at Cys41 (in D-loop), did not show differences in the conformation of the C-terminal segment of F-actin in the presence of different tropomyosins with and without cofilin 1. However, they showed that tropomyosin isoforms differentially regulated cofilin-induced conformational rearrangements at longitudinal and lateral filament interfaces (Ostrowska-Podhorodecka Z *et al.*, 2020). Thus, although the overlap complex’s importance has been well established, it has remained unclear which step in the assembly process is most affected by differences in the N- or C-terminus.

One more vital factor to consider is the binding behaviour of Tpms. As a long lateral size (approximately 30nm) protein, their binding behaviour is described as “Cooperative” (Tobacman, 2008). Thus a numerical approach is used to estimates the binding of Tpms to actin as an explicit function of the free Tpm concentration. The overall affinity (Kapp) of tropomyosin to the linear lattice of the actin filament is dependent on both cooperative binding behaviour and affinity. Other studies suggested that the initial nucleation of tropomyosin molecules on an actin filament is affected by the cooperativity effect. These early assemblies are also indicated to cause structural changes, explaining the sorting of tropomyosin molecules in an isoform-dependent manner and generation of homopolymers (Skórzewski, Śliwińska, *et al.*, 2009a).

Our findings show Tpm1.1 and Tpm1.6 can both nucleate on the same filament while Tpm1.1 always occupied more binding sites on both α and β F-actin than Tpm1.6. We depict that in the absence of a cellular regulatory environment and at the early stages of self-assembly of tropomyosins to the actin filament, most filaments are decorated with more than one type of Tpm and form “heteropolymers”. In the physiological environment, cytoskeletal tropomyosins, in particular Tpm1.6, have been shown to appear as homopolymers (Gimona, Watakabe, and Helfman, 1995; Coulton *et al.*, 2010; Tojkander *et al.*, 2011; Stehn *et al.*, 2013). Interestingly, in our experimental conditions, the tropomyosin concentrations (0.015-0.03 μM) used for imaging experiments were below the range in which the cooperative effect of the binding to actin filaments was observed in our co-sedimentation assays (~0.1 μM). However, in both imaging and bulk biochemistry assays, we observed a higher affinity of Tpm1.1 to actin filaments than Tpm1.6. These findings are in agreement with current models of binding of a large ligand to extended lattices proteins. They also support the notion that the initial stage of binding of long proteins to each other is mainly affinity-dependent (McGhee and Von Hippel, 1974). Together this appears to indicate that affinity seems to be a predominant factor in determining the binding rate of tropomyosin isoforms to actin filaments. Perhaps this is an essential early assembly step before cooperativity initiates.

A recent intravital study on salivary gland acinar cells shows recruitment of Tpms occurs at the same time as, or shortly after, actin assembly (Masedunskas *et al.*, 2018). Similar conclusions are drawn from, the model proposed by Holmes and Lehman (Holmes and Lehman, 2008) and experimental studies by Singh and Hitchcock-DeGregori (Smith and Geeves, 2003; Singh and Hitchcock-Degregori, 2006). Direct visualization of actin filament polymerization and tropomyosin binding at the early assembly is essential to completing the picture.Thus, it appears that the association of tropomyosin to an actin filament can only be explained by considering the whole binding process including, a) the initial affinity dependent interaction of single tropomyosin dimers with the filament, b) the subsequent head-to-tail association of tropomyosins, and c) copolymerization of Tpm and actin. This was captured in the current study with the combination of controlled environment assembly via microfluidics, high spatio-temporal imaging via TIRF microscopy and the computational detection of binding events.

## Acknowledgments

**General**: We thank the Biomedical Imaging Facility of Mark Wainwright Analytical Center at UNSW Sydney for access to microscopes and training facilities. We thank Associate Professor Till Böcking and Professor Peter Gunning for advice and critical feedback on the manuscript.

## Funding

This work was supported by APP1098870 from the NHMRC.

## Author contributions

P.O. and E.P. conceived the idea for this project. P.O and E.P. wrote the manuscript. P.O. and T.S. performed experiments. P.O., P.E. P.N. and A.C. analyzed and interpreted data. All authors discussed the results and critically read the manuscript.

## Competing interests

The authors declare no conflict of interest.

## Data and materials availability

The code used to perform and analyze the data discussed in this paper are available upon request from the corresponding author.

## Supplementary materials

## Introduction

The custom-written single-molecule algorithm developed for quantifying the number of single-molecule of tropomyosins on actin filaments in this study was verified using simulated data. For example, diffraction-limited spots were simulated and recovered from images with varying background noise or located at different distances from one another. These findings showed that our custom-written algorithm has 80% accuracy in estimating the number of single particles in the experimental data.

## Results

The algorithm examines each diffraction-limited spot on a given filament in one iteration. In each iteration, the highest peak is located, analyzed and subtracted from the image. The coding procedure was deciphered when each iteration was outputted into an image. The outcome showed that the diffraction-limited spot with the highest amplitude was initially selected as the first point for analysis, and this process continued until reaching the background signal (Fig. S1A). Further analysis using a simulated image with multiple diffraction-limited spots accumulated in each position demonstrated that the code could recover 1-7 diffraction-limited spots with no error and 8-15 particles with above 88% accuracy (Fig. S1B).

Next, we calculated the signal to noise ratio (SNR) in experimental data as a reference point. We selected random experimental data for the reference image containing images of labeled actin filaments decorated with tropomyosins. The SNR of the first frame in the experimental data for filament images and the single tropomyosin regions were 19.8±3.7 (n=5) and 17±2 (n=7 regions), respectively (Fig. S1C). Subsequently, we investigated the algorithm’s ability to recover diffraction-limited spots in simulated images with varying SNR. Our findings demonstrated 84.4±4% and 90±4% accuracy in recovering a single particle embedded in SNR of 13.3 17.8, respectively (Fig. S1D, left panel). The recovery accuracy in perfect data (i.e. SNR of 55.3) was 93±2% (Fig. S1D, right panel).

Analyzing an elongated diffraction-limited spot requires cutting an ROI of 4×4 pixels around each spot. These occasionally leaves small shoulders behind. The algorithm is equipped with a merging function to marry particles where the centres of the diffraction-limited spots are closer than half of the sigma (sigma is ~1.69 pixels). Furthermore, the precision of the merging function was examined by evaluating the recovery of the spots with varying distances from one another. These data were recovered with 90% accuracy when two diffraction-limited spots were located 3.3 × δ (δ= Gaussian e^-2 radius is set on 1.69 pixels) apart. This value reached 98% accuracy for two diffraction-limited spots with distances equal to and greater than 3.5 × δ (Fig. S1E-F).

Images of diffraction-limited spots representing single particles with known parameters were generated and analyzed to validate the performance of our custom-written algorithm. To determine whether the code can analyze long filaments, we sought to evaluate the recovery of 50 consecutive diffraction-limited spots on one filament. The length of this stimulated filament (472 pixels, ~50 μm) is almost ten times longer than the filaments observed in experimental data (~5 μm). Usually, a filament in experimental data is coated with no more than 25 diffraction-limited spots (referred to as clusters of tropomyosins molecules in the experimental data). Analyzing this long filament, we demonstrated that the algorithm is well equipped to detect information from a long filament (Fig. S2A).

These findings showed that the algorithm is equipped with a merging function to marry close particles belonging to one signal from an elongated diffraction-limited spot.

## Methods

To generate simulated images containing diffraction-limited spots, the “spotmaker” function in MATLAB^®^ was used (written by Tristan Ursell (2012). The images were analyzed with the custom-written single-molecule algorithm.

### Detection of single particles on a long filament

An image containing 50 consecutive diffraction-limited spots was generated. Each diffraction-limited spot has an airy profile with amplitude and sigma of 345.47±0.42 and 1.6902±0.0010, respectively.

### Visualizing the algorithm process

To illustrate the process of the algorithm through each iteration, we simulated an image of nine consecutive diffraction-limited spots with different amplitudes (344±43) and constant sigma (1.69±0.001) values. The algorithm is set to operate by finding the local maximum intensity in the fluorescence image and subsequently fit it with a 2D Gaussian function in its neighbourhood of 4 by 4 pixels. The fitted Gaussian is then subtracted from the image. The process is repeated for the second iteration until the signal level reaches background noise (i.e. the standard deviation of the signal is equal to or smaller than the background noise).

### Detection of the number of single particles per spot in simulated images

The algorithm accuracy in recovering the number of simulated diffracting-limited spots per spot location was evaluated by analyzing images containing nine simulated diffracting-limited spots. Each simulated spot was designed to have an airy pattern with the amplitude and the sigma of 345.4±5, 1.69±0.2, respectively. The background noise of each image was set to 10±5 (a.u.). These values were estimated based on the preliminary observation in the experimental images. Approximately 2-10 diffracting-limited particles (Tpms) were detected in each cluster of proteins on a given filament (data not shown).

### Detection of the single particles in the simulated image with different signal to noise ratio

Recovering simulated diffraction-limited spots embedded in different signal to noise ratios was also tested by simulating 30 consecutive diffraction-limited spots in images with different SNR (image size 40×325 pixels matrix). SNR was set in an image by modulating the “noise-amplitude” parameter in the “Spotmaker” function while keeping the amplitude of diffraction-limited spot constant at 345.45±0 (a.u.). Subsequently, the SNR value in each simulated image was determined by the following equation (1):

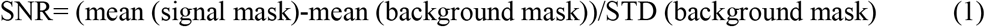

where a rectangular ROI defines the “signal mask” in the middle of the image containing the diffraction-limited spots and “background mask” is the ROI from the whole image excluding the signal mask. For the signal mask, the mean of the 30 peaks was determined by their local-maxima amplitude.

### Merging of the single particles located within a certain distance from each other

To evaluate the merging function in our algorithm, we varied the distance between diffraction-limited spots and set them (Gaussian e^-2 radius) between 1.5 and 3.5 pixels apart. Every single particle has amplitude, sigma and image noise of 345.4±5, 1.69±0.2 and 10±5 (a.u.), respectively. The merging function marries particles where the centres of the diffraction-limited spots are closer than half of the sigma (sigma is ~1.69 pixels).

---

**Figure S1.**
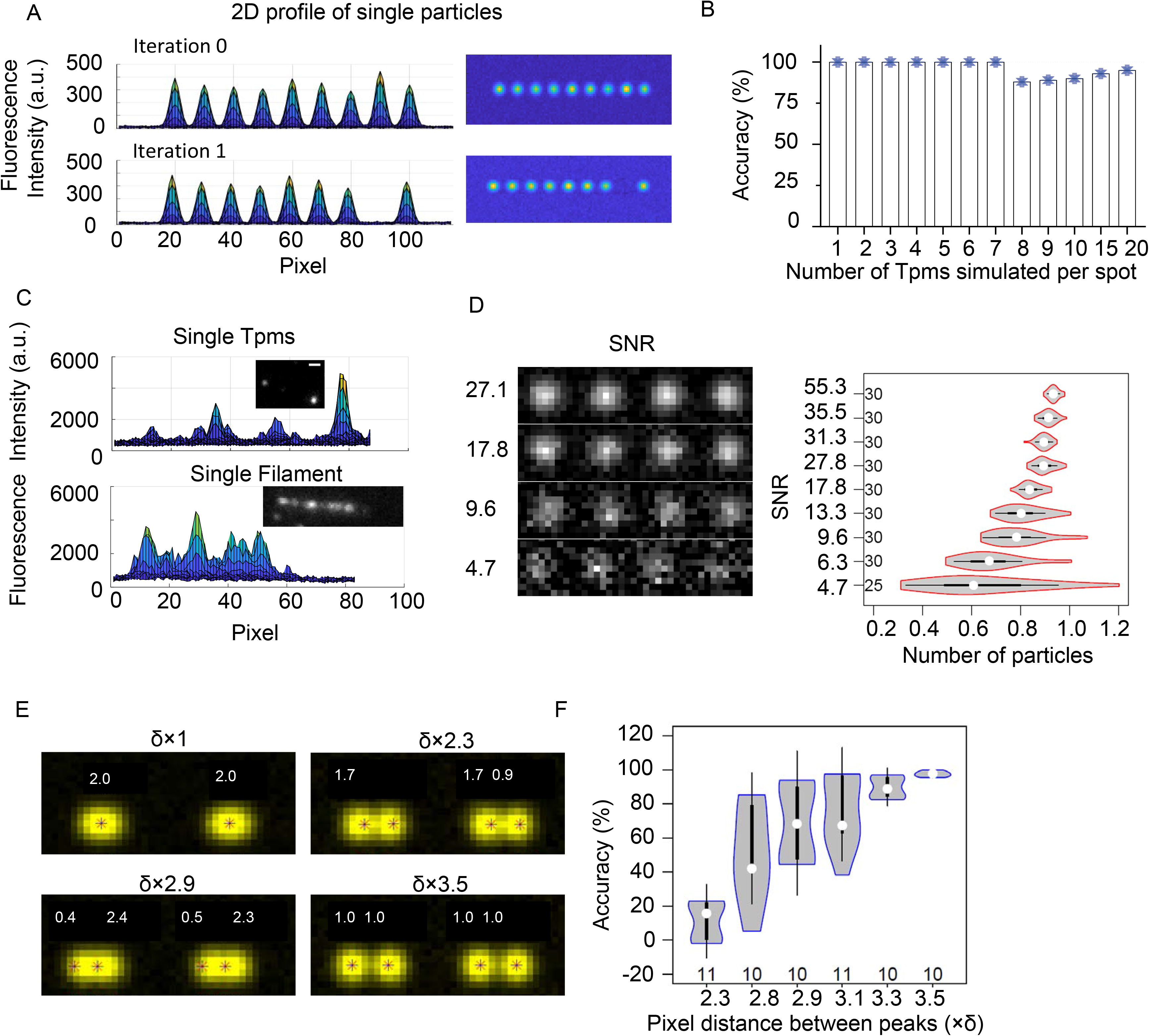
Performance of the algorithm for detection and quantification of the number of molecules in punctate signals along filaments. (A) Iterative detection and depletion of the signal with the highest amplitude in a region of interest. Line profile and a corresponding image containing nine simulated diffraction-limited spots (top) with an amplitude of 344±43 a.u. and a width (σ) of 1.69±0.001 pixels. The first iteration through the algorithm leads to detecting the highest peak and its replacement with average background noise (bottom). (B) Accuracy of the technique for determining the number of molecules from simulated diffraction-limited spots containing 1-20 fluorophores. (C) 2D side-view intensity profiles and corresponding TIRF images (from the experiment) of surface-bound Tpm molecules (top) and Tpm molecules are decorating a surface-bound actin filament (bottom) with a signal-to-noise ratio (SNR) of ~17. (D) Image of simulated diffraction-limited spots with different SNRs (left). The violin plots show the distribution of the number of recovered single particles from a simulated image with 30 spots with a given SNR (inside axes numbers represent the number of peaks). (E) Representative simulated images are depicting sets of two particles with a defined separation (1-3.5 pixel(s)×σ (σ=1.69 pixels). (F) Violin plots showing the distribution of the diffraction-limited spot detection accuracy with different distances from each other (inside axes numbers represent the number of pairs of peaks).

**Figure S2.**
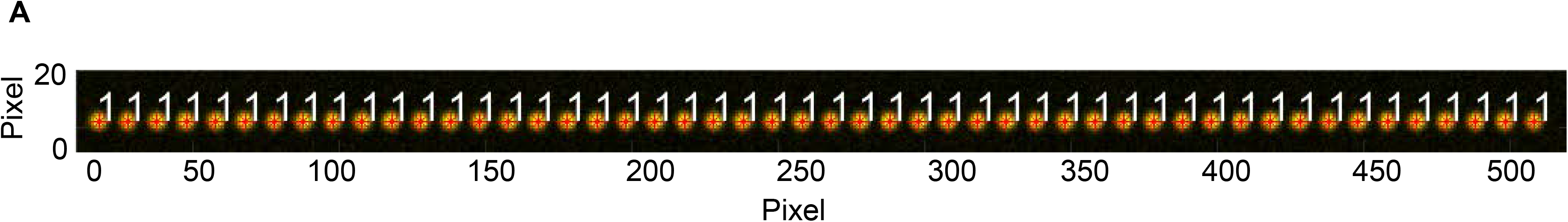
(A) The simulated image contains 50 consecutive diffraction-limited spots (amplitude 345.47±0.42 a.u. and sigma 1.6902±0.0010 pixels). (B) Selection of the diffraction-limited spot with the highest amplitude value displaying the algorithm procedure. (C) Scatter plot with a bar graph of raw data retrieved from 1-20 diffraction-limited spot(s) in each location.

## Notes

### Competing Interest Statement

The authors have declared no competing interest.

